# Gene expression noise produces cell-to-cell heterogeneity in eukaryotic homologous recombination rate

**DOI:** 10.1101/563643

**Authors:** Jian Liu, Jean-Marie François, Jean-Pascal Capp

**Affiliations:** INSA/Université de Toulouse, Laboratoire d’Ingénierie des Systèmes Biologiques et des Procédés, UMR CNRS 5504, UMR INRA 792, Toulouse, France

**Keywords:** Stochastic gene expression, recombination, *Saccharomyces cerevisiae*, yeast, single-cell analysis, rate of evolution

## Abstract

Variation in gene expression among genetically identical individual cells (called gene expression noise) directly contributes to phenotypic diversity. Whether such variation can impact genome stability and lead to variation in genotype remains poorly explored. We addressed this question by investigating whether noise in the expression of genes affecting homologous recombination (HR) activity either directly (*RAD52*) or indirectly (*RAD27*) confers cell-to-cell heterogeneity in HR rate in *Saccharomyces cerevisiae*. Using cell sorting to isolate subpopulations with various expression levels, we show that spontaneous HR rate is highly heterogeneous from cell-to-cell in clonal populations depending on the cellular amount of proteins affecting HR activity. Phleomycin-induced HR is even more heterogeneous, showing that *RAD27* expression noise strongly affects the rate of recombination from cell-to-cell. Strong variations in HR rate between subpopulations are not correlated to strong changes in cell cycle stage. Moreover, this heterogeneity occurs even when simultaneously sorting cells at equal expression level of another gene involved in DNA damage response (*BMH1*) that is upregulated by DNA damage, showing that the initiating DNA damage is not responsible for the observed heterogeneity in HR rate. Thus gene expression noise seems mainly responsible for this phenomenon. Finally, HR rate non-linearly scales with Rad27 levels showing that total amount of HR cannot be explained solely by the time- or population-averaged Rad27 expression. Altogether, our data reveal interplay between heterogeneity at the gene expression and genetic levels in the production of phenotypic diversity with evolutionary consequences from microbial to cancer cell populations.

## 1. Introduction

Expression variations of genes linked to DNA repair and recombination often affect genome stability (Stirling et al., 2011; Ang et al., 2016; Duffy et al., 2016). Whether variable expression levels from cell-to-cell due to gene expression noise could affect homologous recombination (HR) rate and thus genome stability in different subpopulations of clonal populations has not been addressed yet. Noise in gene expression is the variation in the expression level of a gene under constant environmental conditions (Raser and O’Shea, 2005). Downstream effects of noise can have profound phenotypic consequences, drastically affecting gene expression (Blake et al., 2003). This variation in gene expression among genetically identical individual cells could be an advantage in that it would allow heterogeneous phenotypes even in clonal populations, enabling a population of organisms to contain subpopulations with different behaviours and favouring emergence of adapted cells upon environment fluctuation and/or stress conditions (Fraser and Kaern, 2009). Interestingly, genes involved in environmental stress response and metabolism have higher levels of expression noise compared to genes of other biological function in yeast and bacteria (Bar-Even et al., 2006; Newman et al., 2006; Silander et al., 2012). Nevertheless, noise in the expression of precise genes has rarely been shown to be the source of advantageous phenotypic heterogeneity (bet-hedging strategy) and few studies have investigated fitness effects of noise (Viney and Reece, 2013; Liu et al., 2016).

In *S. cerevisiae* expression noise in stress resistance genes confers a benefit in constant stressful conditions because it generates, in the absence of stress, a phenotypic diversity that makes the presence of pre-adapted cells more probable (Blake et al., 2006; Smith et al., 2007; Liu et al., 2015). In addition, recent works showed that heterogeneity in resistance phenotypes due to noise clearly promotes evolvability and shapes mutational effects, partly by modulating the adaptive value of beneficial mutations (Bodi et al., 2017). Also, noise in the expression of genes involved in the DNA replication, repair and recombination processes could directly produce cell-to-cell heterogeneity in the rate of mutation and/or recombination that would also have consequences in terms of evolvability of the population in selective environments (Capp, 2010). Such heterogeneity in mutation rate were recently theoretically studied at various evolutionary timescales (Alexander et al., 2017).

Impact of noise in gene expression on cellular response to DNA damage was investigated in *Escherichia coli* by monitoring the impact of expression variation of the Ada protein in response to DNA alkylation damage (Uphoff et al., 2016). These authors showed that variable induction times of the damage response were observed depending on the initial expression level of Ada, with cells that do not respond for generations because no Ada proteins are initially expressed. This creates a subpopulation of cells with an accumulation of foci of the DNA mismatch recognition protein MutS used as a marker for labeling nascent mutations (Uphoff et al., 2016), showing heterogeneity in the mutation rate at the single-cell level. The conclusion of the study highlighted that non-genetic variation in protein abundances thus leads to genetic heterogeneity. Nevertheless, this measurement remains an indirect evaluation of the genetic heterogeneity through the detection of a mismatches biosensor. Moreover neither the genetic consequences of noise in expression of DNA repair genes on a genomic substrate, nor its subsequent phenotypic consequences, were analyzed. Finally investigating similar phenomena in eukaryotes is motivated by their higher number of different proteins and more complex pathways involved in the DNA replication, repair and recombination processes that diversify and multiply the possible sources of cell-to-cell variation in mutation and recombination rate.

For simplicity, HR can be defined as the repair of DNA lesions based on homologous sequences (Symington et al., 2014). It underlies a number of important DNA processes that act to both stabilize (e.g. repair of DNA double-strand breaks (DSBs)) and diversify (generation of crossover during meiosis) a genome. Meiotic HR rate for instance has revealed considerable inter-individual differences (Dumont et al., 2009) or extensive variations along chromosomes (Kauppi et al., 2004). But technical limitations only allowed studies on whole cell populations, providing an averaged view of this process. Only recent studies of meiotic HR have revealed the diversity in crossover frequency in single sperm cells (Lu et al., 2012; Wang et al., 2012) or oocytes (Hou et al., 2013). Spontaneous mitotic HR rate also varies along chromosomes, with for instance elevated recombination rates in transcriptionally active DNA (Thomas and Rothstein, 1989), but analysis of cell-to-cell heterogeneity in mitotic HR rate in clonal cell populations is still lacking.

Mitotic HR is entirely conservative when it occurs following DNA replication where a sister chromatid is available as a template. However, HR acting on DSB can produce genome instability, especially when utilizing sequences on a homologous chromosome that can lead to crossovers and potential loss of heterozygosity, or when occurring between dispersed repeated DNA. Indeed interrepeat recombination can cause deletions, duplications, inversions or translocations, depending on the configuration and orientation of the repeat units. These non-conservative events are especially studied in this work because there are of major importance for evolution.

HR pathways are particularly well-documented in *S. cerevisiae* (Paques and Haber, 1999; Symington et al., 2014). A diversity of mechanisms can modify HR activity, either indirectly by increasing the generation of DNA lesions, or directly by blocking the completion of HR and/or altering the kinetics of genetic recombination and the assembly/disassembly of the HR protein complexes (Alvaro et al., 2007). Each class of mechanism is respectively well-represented in *S. cerevisiae* by the absence of the *RAD27* and *RAD52* genes. On one hand, Rad52 is involved in multiple pathways of repairing DSB (Symington, 2002). It binds single-stranded DNA to stimulate DNA annealing and to enhance Rad51-catalyzed strand invasion during the HR process called synthesis-dependent strand annealing (New et al., 1998; Song and Sung, 2000). It is also involved in Rad51-independent pathways used to repair DSB such as single-strand annealing (SSA) (Symington et al., 2014). SSA is stimulated if the DSB lies in a unique sequence between two repeated sequences and can lead to the repeat contraction or expansion. The various roles of Rad52 explain the highly defective mitotic recombination in *rad52 S. cerevisiae* mutants (Dornfeld and Livingston, 1992; Rattray and Symington, 1994). On the other hand, the *RAD27* gene of *S. cerevisiae* encodes a 5′-3′ flap exo/endonuclease, which is a functional homolog of mammalian FEN1/DNaseIV that plays an important role during DNA replication for Okazaki fragment maturation (Balakrishnan and Bambara, 2013). It cleaves the unannealed 5′ “flap” structure containing the primer that appears in 5’ of the previous Okazaki fragment after synthesis of the next one (Zheng and Shen, 2011). The absence of *RAD27* generates an accumulation of 5’ flap structures that can be resolved by the Rad52 dependent-HR pathway (Debrauwere et al., 2001). *S. cerevisiae* rad27Δ mutants accumulate of single- and DSB (Tishkoff et al., 1997) and display a broad array of defects in genome stability including an increased spontaneous recombination (Johnson et al., 1995; Sommers et al., 1995; Tishkoff et al., 1997).

Here we choose to use a *S. cerevisiae* strain containing a HR substrate that allows measuring the rate of non-conservative interrepeat recombination events and to sort subpopulations depending on the native expression level of *RAD27* and *RAD52*. The antagonist effects of their deletion on HR frequency and their different modes of action (direct or indirect) to affect HR led us to choose these two genes to study the influence of their heterogeneous cellular amounts. This study provides evidence that cell-to-cell expression fluctuations of Rad27 and Rad52 produce heterogeneity in both spontaneous and induced HR frequency in the population. Moreover HR rate non-linearly scales with Rad27 levels. The recombination rate varies strongly above the mean Rad27 expression level of the population before reaching a plateau at its highest values for the highest expression levels. Finally, it does not result from differences in cell cycle distribution, and can be hardly explained by heterogeneity in DNA damage because it occurs also when cells are simultaneously sorted at equal level of the Bmh1 protein that is upregulated by DNA damage. Altogether, these results showed that noise in the expression of genes involved in DNA transactions can lead to heterogeneous homologous recombination rate between individual eukaryotic cells.

## 2. Material and Methods

### 2.1 Yeast strains and growth conditions

All the strains and primers used in this work are listed in Supplementary Tables 2 and 3, respectively. The strain KV133 (Verstrepen et al., 2005) (BY4742 *MATα; his3*Δ*1; leu2*Δ*0; lys2*Δ*0; ura3*Δ*0 FLO1::URA3*) (URA3 inserted in the middle of the tandem repeats) was kindly provided by Kevin J Verstrepen (KU Leuven). To create strain JA0200 from KV133, a PCR fragment containing *LEU2* and its native promoter and terminator was amplified from the genomic DNA of the S288c strain with primers F1 and R1, and transformed into KV133. The construction was verified by PCR with primers C1 and C2. To create the strains containing the fusion *RAD27-YFP* and *RAD52-YFP* (JA0219 and JA0220 respectively), PCR fragments containing *YFP-kanR* and homologies to *RAD27* or *RAD52* were amplified with primers F2 and R2, or primers F3 and R3 respectively, from the plasmid pfa6a-*YFP-kanR* (constructed in our lab), and transformed into JA0200. The constructions were verified by PCR with primers C3 and C4, or C5 and C4 respectively. To create the strains containing the double fusion *RAD27-YFP-tdTomato* and *RAD52-YFP-tdTomato* (JA0240 and JA0241 respectively), PCR fragments containing *tdTomato-SpHis5* and homologies to *YFP* were amplified with primers F4 and R4 from the plasmid pfa6a-link-*tdTomato-SpHis5* (Addgene), and transformed into JA0219 and JA0220. The constructions were verified by PCR with primers C3 and C6, or C5 and C6 respectively. To delete *RAD27* (strain JA0217), a PCR fragment containing *LYS2* and homologies to *RAD27* was amplified from the genomic DNA of the S288c strain with primers F6 and R6, and transformed into JA0200. The construction was verified by PCR with primers C7 and C8. To insert *pBMH1-yEGFP* into the strains JA0240 and JA0200 (strains JA0242 and JA0243 respectively), the integrative plasmid pJRL2-*pBMH1-yEGFP* containing homologies to *LEU2* was cut by AscI (New England Biolabs) and transformed. The construction was verified by PCR with primers C9 and C2. All the transformations were carried out by the standard lithium acetate method.

All the strains were grown in liquid YNB medium (20 g/L glucose (Sigma), 1.71 g.L^−1^ yeast nitrogen base without amino acids and nitrogen (Euromedex) and 5 g.L^−1^ ammonium sulfate (Sigma)) at 30°C with rigorous shaking (200 rpm). Auxotrophic strains were supplemented with the required molecules at the following concentrations: 0.02 g.L^−1^ histidine (Sigma), 0.05 g.L^−1^ lysine (Sigma) and 0.1 g.L^−1^ leucine (Sigma). For phleomycin treatment, cells in stationary phase were diluted 100 times in YNB medium containing 5 µ g.mL^−1^ phleomycin (Sigma) and grown at 30°C with rigorous shaking (200rpm) for 16 hours.

The YPD plates contained 20 g.L^−1^ glucose, 20 g.L^−1^ agar (Euromedex), 10 g.L^−1^ peptone (Euromedex) and 10 g.L^−1^ yeast extraction (Euromedex). The 5-FOA and CAN plates contained 20 g.L^−1^ glucose, 20 g.L^−1^ agar, 1.71 g.L^−1^ yeast nitrogen base, 5 g.L^−1^ ammonium sulfate, 0.79 g.L^−1^ complete supplement mixture (Euromedex) and 1 g.L^−1^ 5-FOA (Euromedex) or 0.06 g.L^−1^ canavanine (Sigma) respectively.

### 2.2 Fluorescence activated cell sorting

The cell sorting experiments were carried out on the MoFlo Astrios EQ cell sorter with the Summit v6.3 software (Beckman Coulter). Cells in stationary phase were diluted 100 times and grown at 30°C with rigorous shaking (200 rpm) for 16 hours prior to cell sorting (final OD ≈ 2). Cultures were spun down at 3000g for five minutes at 4°C. Growth media was removed and cells were re-suspended in ice cold PBS. The SmartSampler and CyClone tubes holder were kept at 4°C during cell sorting. Cell sorting was carried out with 70µm nozzle and 60psi operating pressure. The sorting speed was kept around 30 000 events per second. The purity mode for the sort mode and 1 drop for the droplet envelope were chosen. Based on the FSC-Area *vs* SSC-Area (488 nm laser) plot and the FSC-Height *vs* FSC-Area (488 nm laser) plot, single cells with similar cell size and granularity were first selected. Then based on the histogram of the YFP-tdTomato fluorescence (560 nm laser, 614/20 filter), single cells with 10% highest fluorescence and 10% lowest fluorescence were sorted simultaneously (Figure 2); or single cells were sorted simultaneously into five subpopulations distributed as follows: 0-10%, median between the median of the 0-10% and the median of the whole population +/- 5%, median of the whole population +/- 5%, median between the median of the whole population and the median of the 90-100% +/- 5%, 90-100% (Figure 3). This division allowed reproducible sorting between replicates even if slight variations of the distribution of absolute expression levels occurred.

**Figure 1.**
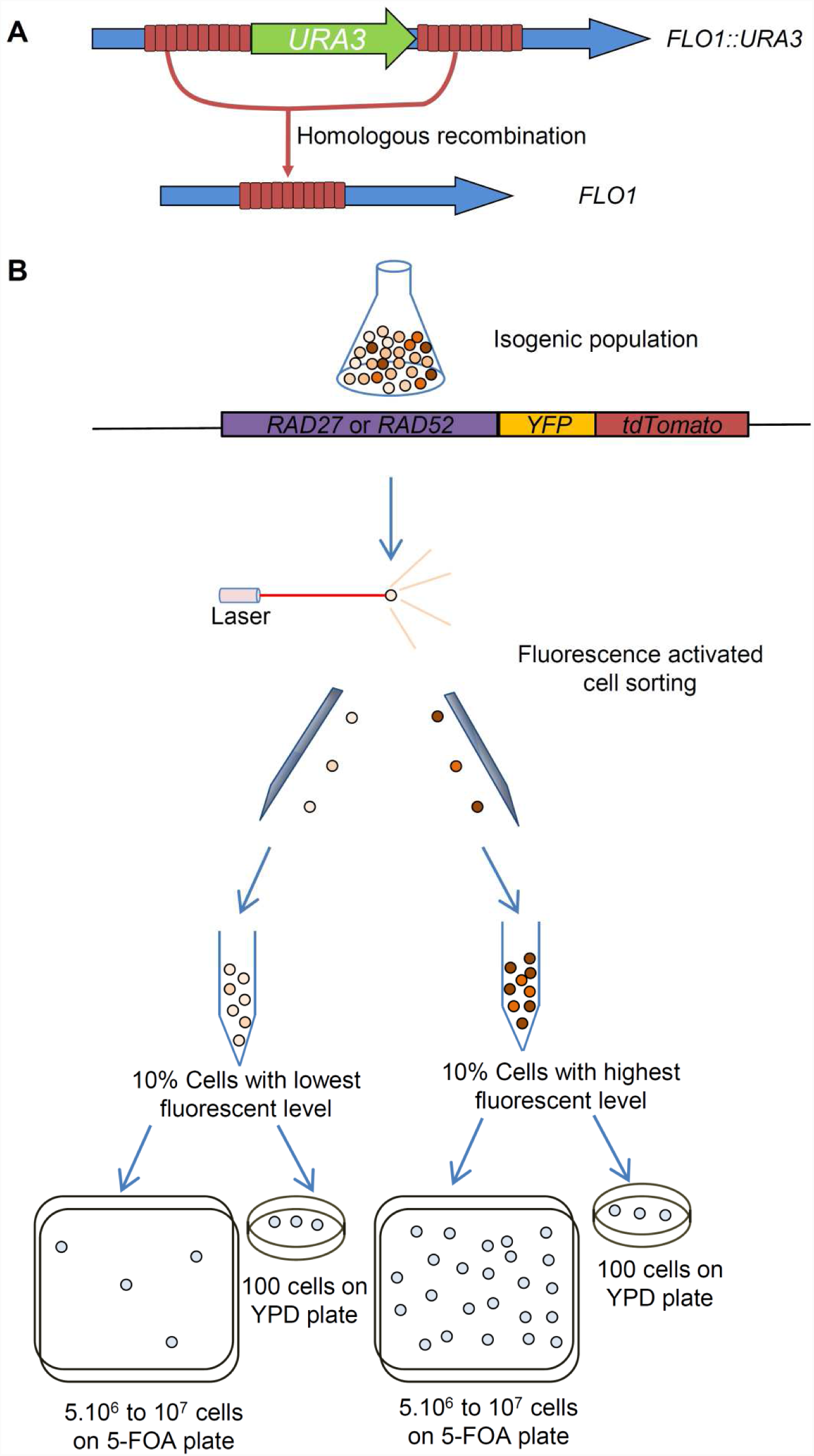
Experimental procedure. (A) Homologous recombination (HR) frequency between intragenic repeats in *FLO1* was measured by the loss of the *URA3* expression cassette integrated in the middle of the *FLO1* repeats (Verstrepen et al., 2005). When a recombination event occurs in the repeats, the *URA3* marker loss results in a 5-FOA resistant (Ura^−^) strain containing a new *FLO1* allele. (B) The double fluorescent marker YFP-tdTomato was fused to either *RAD52* or *RAD27* at their original genomic locus in the strain harbouring the recombination substrate, allowing sorting of cells with extreme expression levels. 5.10^6^ to 10^7^ cells were sorted for each subpopulation, and spread on 5-FOA plates. In parallel viability was evaluated on YPD plates, allowing calculation their respective rate of loss of *URA3* function.

**Figure 2.**
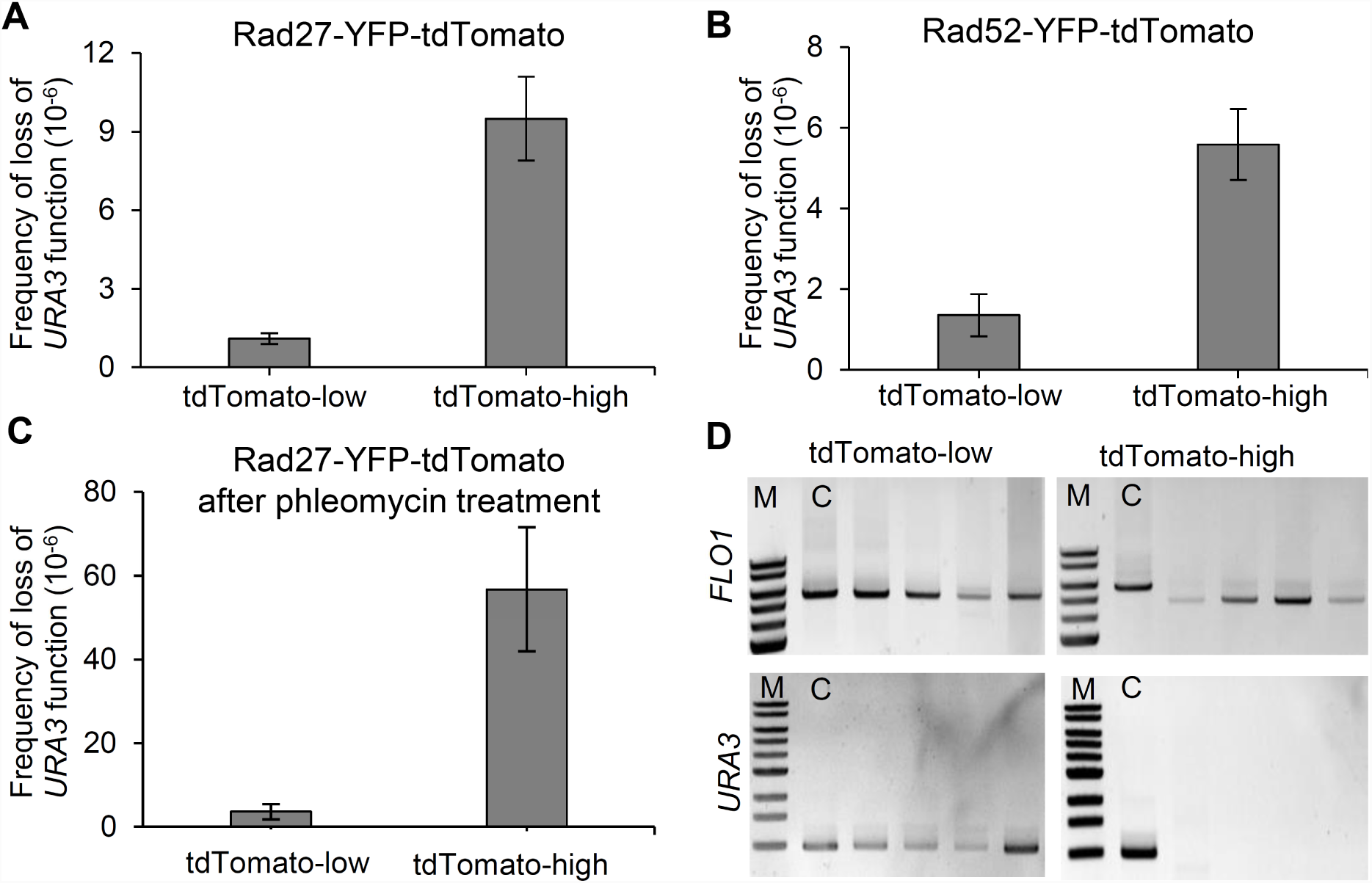
Noise in the expression of genes affecting HR activity produces cell-to-cell heterogeneity in spontaneous HR rate. (A) Spontaneous frequency of loss of *URA3* function in the subpopulations with the highest (10%) and lowest (10%) Rad27-YFP-tdTomato cellular amounts. (B) Spontaneous frequency of loss of *URA3* function in the subpopulations with the highest (10%) and lowest (10%) Rad52-YFP-tdTomato cellular amounts. (C) Phleomycin-induced frequency of loss of *URA3* function in the subpopulations with the highest (10%) and lowest (10%) Rad27-YFP-tdTomato cellular amounts. Results are the mean of 3 independent experiments with standard deviation. (D) Examples of PCR amplification of the new *FLO1* alleles in 5-FOA resistant clones showing that their length is modified in the high-expressing subpopulations, and not in the low-expressing subpopulations compared to the control strain (C). PCR amplification of the *URA3* gene in the same clones showed that it is lost by HR in the high-expressing subpopulations and still present in the low-expressing subpopulations.

**Figure 3.**
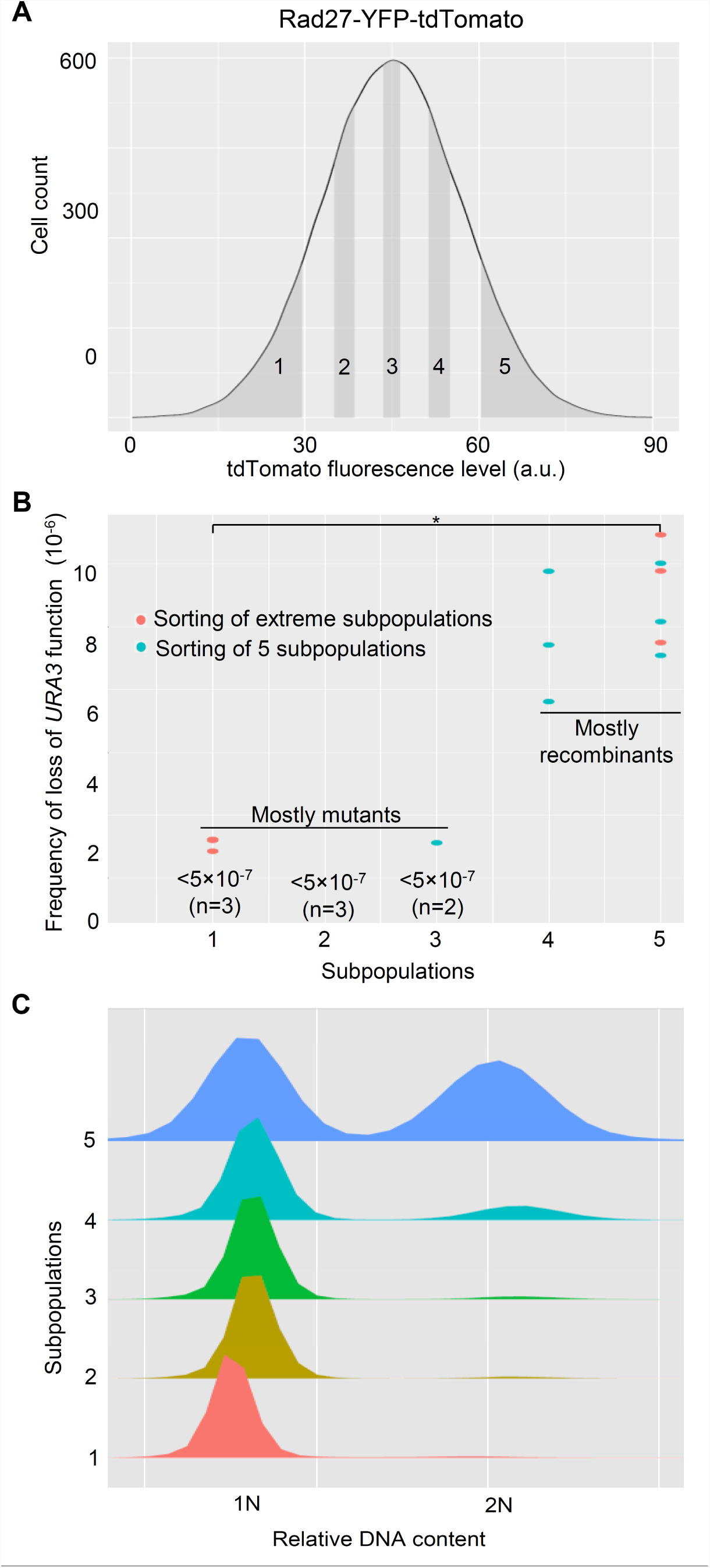
HR rate non-linearly scales with Rad27 levels and is weakly correlated with differences in cell cycle distribution. (A) Five subpopulations homogenously distributed in the whole population were sorted thanks to the fused protein Rad27-YFP-tdTomato. Each subpopulation represents 10% of the whole population. They are numbered 1 to 5 from the lowest to the highest expression levels. About 2.10^6^ cells were sorted for each subpopulation (3 independent experiments), and spread on 5-FOA plates. In parallel viability was evaluated on YPD plates, allowing calculation their respective frequency of loss of *URA3* function. 3 independent experiments were performed. (B) Measurable rates on these five subpopulations (in blue) were combined to the results obtained in Figure 2A (in red) to plot the relationship between rate of loss of *URA3* function and Rad27 levels. Each dot represents one sorting experiment for one subpopulation that has given a measurable rate. When no rate was measurable because of the absence of 5-FOA resistance clone, the maximal rate is written. As shown in Figure 2D, 5-FOA resistance is due to mutation-based inactivation of the *URA3* gene in subpopulations1 to 3 and to recombination–based loss of the *URA3* marker in subpopulations 4 and 5. A significant statistical difference is represented by (*) when p<0.05 in Wilcoxon signed rank test. (C) Cell cycle distribution in the five subpopulations isolated from the Rad27-YFP-tdTomato-expressing population is represented.

To sort GFP and YFP-tdTomato simultaneously, the fluorescence of the strains with only GFP (488 nm laser, 526/52 filter, strain JA0243) or YFP-tdTomato (560 nm laser, 614/20 filter, strain JA0240) was first measured. There is only negligible overlap between these fluorophores, hence there was no need for compensation. Then based on the GFP *vs* YFP-tdTomato plot of the strain JA0242, 5% single cells of the total population with similar GFP fluorescence as the mean of the population but extreme YFP-tdTomato fluorescence were sorted, as well as 5% single cells of the total population with similar YFP-tdTomato fluorescence as the mean of the population but extreme GFP fluorescence.

### 2.3 Measurement of HR frequency

To measure the HR frequency of the whole population, 500 µL culture (OD ≈ 2) was spread on 5-FOA petri plates (100×15 mm, Fisherbrand). The culture was diluted 10 000 times and 20 µL diluted culture was spread on YPD petri plates. The plates were kept in 30°C incubator for 3 days and the number of clones was counted. The size of the new *FLO1* alleles from the clones isolated on the 5-FOA plates was analyzed by PCR using primers F5 and R5. The presence of *URA3* was verified using primers C10 and C11. The size of *FLO5* and *FLO9* were further analyzed by primers C13 and R5 or C12 and R5 respectively. Then the frequency of loss of *URA3* function was calculated as follow:

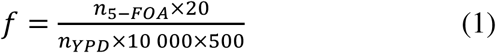

where *f* denotes the frequency of loss of *URA3* function, *n*_5-*FOA*_ denotes the number of clones on the 5-FOA plates, and *n*_*YPD*_ denotes the number of clones on the YPD plates.

To measure the frequency of loss of *URA3* function of the subpopulations, 5.10^6^ to 10^7^ cells (depending on the replicate) of each subpopulation were sorted (around 10 mL) and spread on 5-FOA square culture dishes (224×224×25 mm, Corning). Then 100 cells were sorted and spread on YPD plates. The dishes were kept in 30°C incubator for 3 days and the number of clones was counted. The size of the new *FLO1* alleles, the presence of *URA3* or the size of the *FLO5* and *FLO9* alleles were analyzed by PCR from the clones isolated on the 5-FOA plates using primers F5 and R5, C13 and R5 or C12 and R5 respectively. Then the frequency of loss of *URA3* function was calculated as follow:

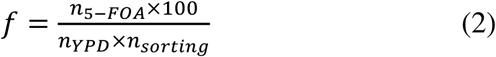

where *f* denotes the frequency of loss of *URA3* function, *n*_5-*FOA*_ denotes the number of clones on the 5-FOA dishes, *n*_*YPD*_ denotes the number of clones on the YPD plates, and *n*_*sorting*_ denotes the number of cells sorted.

### 2.4 Analysis of cell cycle stage distribution

10^6^ cells were sorted and fixed in 70% ethanol at 4°C for at least 12 hours. They were then washed in 50 mM sodium citrate (Sigma) buffer (pH 7.5) and treated by RNAse A (Eurogentec) and proteinase K (Eurogentec). Yo-Pro-I (Thermer Fisher) was used to stain the genomic DNA. The relative DNA content was measured by MACSQuant® VYB flow cytometry (Miltenyi Biotec).

### 2.5 Statistics

The Wilcoxon signed rank test was performed in R (version 3.4.1) with the wilcox.test fonction.

## 3. Results

### 3.1 Noise in the expression of *RAD52* and *RAD27* produces heterogeneity in spontaneous and induced HR rate

The system developed by Verstrepen *et al* to measure non-conservative HR between intragenic tandem repeats (Verstrepen et al., 2005) used the auxotrophic marker *URA3* integrated in the tandem repeats of the *FLO1* gene in *S. cerevisiae* (Figure 1A). As recombinants do not grow on the initial medium where *URA3* is needed for growth because uracil is lacking (no clonal expansion possible), the frequency of yeast cells then growing on 5-FOA-containing medium provides a quantitative estimate of the actual recombination rates (number of events per cell division) as previously suggested (Verstrepen et al., 2005). However the loss of *URA3* function could also arise by direct mutations in the *URA3* coding region.

This system confirmed that the absence of *RAD27* and *RAD52* respectively strongly increases and decreases the HR frequency (Verstrepen et al., 2005). The increased recombination frequency in rad27Δ mutants suggests that *FLO1* repeat instability is associated with the occurrence of DSB due to defective DNA replication (Kokoska et al., 1998). The absence of an effect in rad51Δ mutants and the decrease in recombination observed in various other mutants, especially rad50Δ and rad52Δ, suggests that recombination in this system does not require strand invasion and depends on DSB repair by SSA (Verstrepen et al., 2005). Nevertheless, one cannot exclude that loss of *URA3* could happen preferentially through gene conversion-associated crossing over in wild-type cells and that repair could switch to SSA in rad51Δ mutants.

Subpopulations were sorted based on the expression level of *RAD52* or *RAD27* fused to *YFP* and *tdTomato* at their original genomic locus (Figure 1B). A fluorescent signal above the auto-fluorescence level for the whole population was needed to efficiently sort even the cells expressing *RAD52* and *RAD27* at the lowest levels. Thus, we chose tdTomato which is one the brightest fluorescence protein to be fused to Rad52 and Rad27 because of the low expression of the corresponding genes, and added YFP that improved the fluorescence of our tagged proteins compared to tdTomato only. This YFP-tdTomato double fusion to the C-terminal domain of Rad52 seems not to affect its functionality because the average HR frequency in the population was the same as in the wild-type (Supplementary Figure 1). On the contrary the Rad27-YFP-tdTomato fusion slightly decreased this average HR frequency (Supplementary Figure 1). Nevertheless the functionality of the fused Rad27-YFP-tdTomato protein is close to the native protein because it confers an HR rate that is in the same order of magnitude as the wild-type when compared to the strongly increased HR frequency in rad27Δ mutants (Supplementary Figure 1).

Among the heterogeneous expression levels of these genes at the single-cell level, we first isolated the two extreme subpopulations in terms of fluorescence intensity, each of them representing 10% of the whole population. While viability was similar for both subpopulations (Supplementary Figure 2), the rate of loss of *URA3* function as determined by the frequency of cells growing on 5-FOA plates (Figure 1B) was 10-times higher for the Rad27-high subpopulation (Figure 2A and Supplementary Table 1) and 4-times higher for the Rad52-high subpopulation (Figure 2B and Supplementary Table 1) compared to the low-subpopulations.

PCR amplification of the new *FLO1* alleles in 5-FOA resistant clones showed that *FLO1* is modified in the Rad27-high and Rad52-high subpopulations, and not in the Rad27-low and Rad52-low subpopulations, suggesting that recombination events and rearrangements among tandem repeats indeed led to the loss of *URA3* only in the formers (Figure 2D and Supplementary Figure 3). As 5-FOA resistance can also arise through mutations in the *URA3* coding region, we wanted to actually confirm the nature of the genetic changes by testing the presence of the *URA3* gene in the resistant clones. We confirmed that the loss of *URA3* gene in the Rad27-high and Rad52-high subpopulations occurred by recombination between intragenic repeats in *FLO1* (Figure 2D and Supplementary Figure 3). The variability of the size of the new FLO1 alleles seen in Supplementary Figure 3 shows that these clones likely occurred during independent events and that they were not the result of clonal expansion. On the contrary, 5-FOA resistant clones from the Rad27-low and Rad52-low subpopulations still contained the *URA3* gene at the expected size, showing that they likely acquired 5-FOA resistance by mutation (Figure 2D and Supplementary Figure 3). Thus, the precise difference in HR rate between these extreme subpopulations cannot be quantified because of the absence of detectable recombinant cells in the subpopulations with the lowest expression levels.

The finding that Rad52-high cells harbor higher HR rate is in accordance with its direct involvement in HR pathways. In contrast, it is at first glance counterintuitive to find Rad27-high cells with the highest HR rate considering that the deletion of this gene leads to increased recombination, even if, as mentioned above, Rad27-YFP-tdTomato did not fully recapitulate the functionality of the native Rad27 protein.

To confirm the heterogeneity in spontaneous HR rate produced by the *RAD27* heterogeneous expression levels, we induced the production of DSB by pretreating cells with 5 µg.ml^−1^ phleomycin for 16 h. This chemical is a water-soluble antibiotic of the bleomycin family that catalyzes DSB in DNA (Moore, 1988), thus strongly increasing recombination frequency. Sublethal concentration was used to avoid loss of cell viability in the subpopulations (Supplementary Figure 2) and to affect as little as possible growth (Liu et al., 2015), yet allowing measurable effect of the amount of induced DSB without toxicity. Frequency of loss of *URA3* function is far more induced in the high-subpopulation than in the low-subpopulation following this pretreatment (Figure 2C and Supplementary Table 1). This stronger effect of the drug on the high level population is coherent with the fact that an increased level of Rad27 increases HR.

As it would appear difficult to reliably conclude the general effect of single-cell protein levels on recombination, we also assayed repeat expansion / contraction at other loci (*FLO5* and *FLO9*) in these clones as previously performed among *S. cerevisiae* strains (Verstrepen et al., 2005). No variation was detected (Supplementary Figure 4) but the probability to observe HR events in both of these loci in the same cell during the course of our experiments is extremely low. Even on the whole sorted subpopulations; the rarity of such events makes their detection impossible without any selective pressure to enrich them.

**Figure 4.**
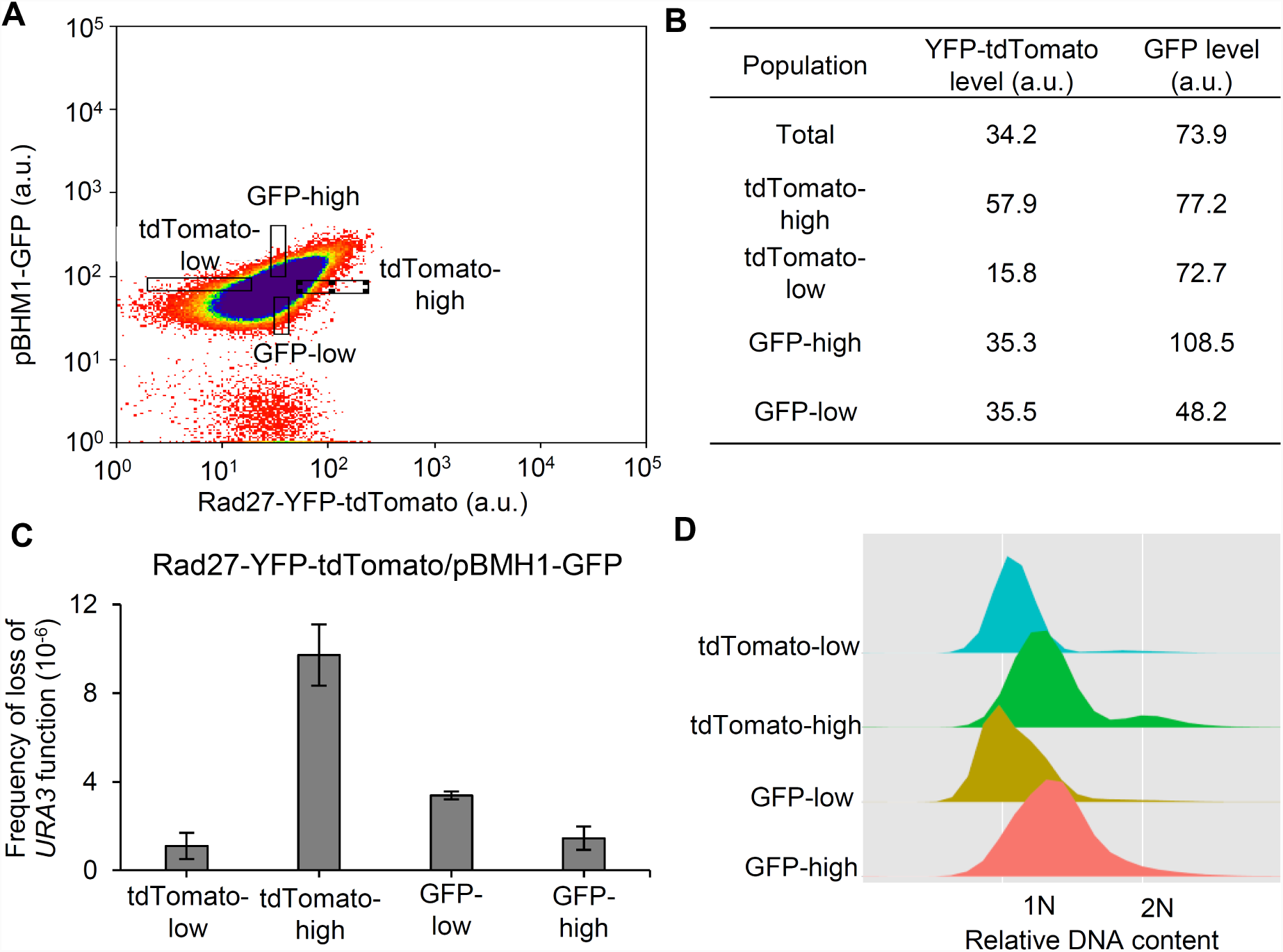
The initiating DNA damage is not responsible for the observed heterogeneity in HR rate. (A) Dot plot of the population expressing *pBMH1-GFP* and *RAD27-YFP-TdTomato*, with gates allowing sorting of cells with similar expression levels of one fluorescent marker and extreme expression levels of the other. (B) Rad27-YFP-tdTomato and GFP levels in the four subpopulations isolated from the previous dot plot. (C) Spontaneous HR frequency in the subpopulations with similar *pBMH1-GFP* expression levels and the highest (10%) and lowest (10%) Rad27-YFP-tdTomato cellular amounts, and in the subpopulations with similar *RAD27-YFP-TdTomato* expression levels and the highest (10%) and lowest (10%) GFP cellular amounts. Results are the mean of 3 independent experiments with standard deviation. (D) Cell cycle distribution in the four subpopulations isolated from the previous dot plot.

### 3.2 HR rate non-linearly scales with Rad27 levels

Considering the high heterogeneity in HR activity observed between Rad27-high and Rad27-low subpopulations, we ask whether the HR rate scales linearly or non-linearly with Rad27 levels. To do so, we performed the same experiment as described in Figure 1B but we sorted cells into five subpopulations homogenously distributed in the population from the lowest (subpopulation 1) to the highest (subpopulation 5) expression levels (Figure 3A). Each subpopulation represents 10% of the whole population. The sorting process is less efficient when cells are sorted into 5 subpopulations instead of two. Far more time is needed to get the same number of cells in a given subpopulation when sorting into 5 subpopulations. Thus we were only able to sort out only around 2.10^6^ for each subpopulation which lasted at least 3 hours. Extending further the duration of the experiment would lead to bias linked to prolonged time in tubes before and after passage into the sorter, to the diluted medium in the harvest tubes… Moreover inducing DSB was not chosen for this experiment because phleomycin treatment slightly increases the *RAD27* expression level so that it does not allow studying basal expression level and spontaneous events that can be considered as more relevant to evolution.

By doing so, we did not detect any 5-FOA resistant cells in subpopulations 1 and 2 in any replicate (Supplementary Table 1). Only one clone was detected in subpopulation 3 in one experiment while many more and similar numbers of clones were observed in subpopulations 4 and 5 (Supplementary Table 1). We confirmed again that these resistant clones are generated by recombination when Rad27 levels are high. As results were similar for subpopulation 5 in both sets of experiments, we chose to combine the results on these five subpopulations with the results from Figure 2A to plot the relationship between the frequency of loss of *URA3* function and Rad27 levels (Figure 3B). We indicated in this plot that resistance is mostly due to mutation-based inactivation of the *URA3* gene in the first subpopulations and to recombination–based loss of the *URA3* marker in subpopulations 4 and 5. Thus, even we were not able to measure HR rate in the formers, it appears that HR rate non-linearly scales with Rad27 levels because it reaches a plateau in the latters after an abrupt variation that occurs at least between subpopulations 3 and 4 slightly above the mean expression level of the population.

Finally, we took advantage of having six independent replicates for high- and low-Rad27 subpopulations to perform a non-parametric statistical test. These data show a significant difference (p=0.03) (Figure 3B), suggesting that for other data where the same tendency is observed with only three replicates (high-vs low-Rad52, high-vs low-Rad27 with phleomycin), similar conclusions could be drawn (these data constitute too small data samples to robustly apply proper statistical analysis).

### 3.3. Differences in cell cycle distribution do not explain heterogeneous HR rate

Since cell-cycle dependence of transcription dominates noise in gene expression (Zopf et al., 2013) and that transcript level of *RAD27* is not constant through the cell cycle (Skotheim et al., 2008), we determined whether the difference in recombination rate in the sorted subpopulations is not a consequence of different cell cycle states. We measured the cell cycle distribution among the same five sorted subpopulations. As the chosen sorting mode on the cytometer excluded most of the budding cells, strong enrichment in G1 cells was expected, that should limit the impact of cell cycle state or size differences. Indeed we observed almost exclusively G1 cells in subpopulations 1 to 3 (Figure 3C). Only subpopulation 5 contained more G2 cells.

This observation suggests that Rad27 expression heterogeneity is not mostly due to different cell cycle states, because the strong increase seen between subpopulations 3 and 4 is associated to only a slight difference in cell cycle distribution exist with a small G2 bump. If this appearance of G2 cells in subpopulation 4 was responsible for the strong increase in recombination frequency, we would expect an even higher increase in subpopulation 5 where G2 are far more abundant. Instead, subpopulation 5 with the highest expression levels harbored the same frequency of loss of *URA3* function as subpopulation 4 in spite of its strong enrichment in G2 cells, making us thinking that cell cycle distribution only poorly influences recombination rate.

### 3.4. Heterogeneity in HR rate does not result from heterogeneity in DNA damage

Given the poor contribution of cell cycle and the unexpected correlation between the *RAD27* expression level and HR rate, we went further in deciphering the origins of the heterogeneity in HR rate. Apart from cell cycle, heterogeneity in DNA damage could affect recombination activity. To study this hypothesis, we performed a double sorting of extreme subpopulations based on the expression of Rad27-YFP-tdTomato on one hand, and the expression of GFP driven by the promoter of the *BMH1* gene (*pBMH1*) on the other hand (Figure 4A and 4B). Bmh1 is one of the two yeast 14-3-3 proteins and many studies also showed the important role of 14-3-3 proteins in DNA duplication and DNA damage response in fungi (Kumar, 2017). Especially, it directly modulates DNA damage-dependent functions of Rad53 (Usui and Petrini, 2007) and it is upregulated by DNA damage along with other protein factors associated with DNA damage response (Kim et al., 2011). Thus sorting cells with extreme levels of Rad27 and simultaneously at equal level of *pBMH1*-driven GFP should ensure observing the phenomenon in cells with similar levels of DNA damage and showing that heterogeneity in DNA damage is not responsible for it. Moreover, it possesses a relatively strong promoter that allows GFP expression largely above the auto-fluorescence threshold.

Frequency of loss of *URA3* function was again higher in Rad27-high than in Rad27-low cells when also sorting cells with the same GFP level (Figure 4C and Supplementary Table 1), with a 10-fold factor similar to the previous experiments (Figure 2A). No difference in viability was observed (Supplementary Figure 5). 5-FOA resistance was again due to recombination in Rad27-high sorted cells and to mutation in Rad27-low sorted cells, showing that the difference is due to differences in HR rate. On the contrary, both GFP-low and GFP-high subpopulations exhibited close frequency of loss of *URA3* function when also sorted at equal level of Rad27. However, we noticed that 5-FOA resistant clones were slightly more frequent in GFP-low cells (Figure 4C). This higher HR rate might be explained by the fact that cells expressing *BMH1* at lower levels might accumulate more DSB, which in turns could slightly enhance HR and/or mutation rate (Engels et al., 2011). Finally when analyzing cell cycle distribution, DNA content plot is shifted to the right for both high GFP and high tdTomato expressing cells (Figure 4D). If cell cycle distribution had a strong influence on recombination rate, both subpopulations would harbour increased frequency. Nevertheless a strong increase in the frequency of loss of *URA3* function is only observed in tdTomato-high cells and not in GFP-high cells (it is even lower in the latter case). This argues again against its contribution in the generation of HR rate heterogeneity.

## 4. Discussion

We observed heterogeneous HR rates in the subpopulations expressing *RAD52* or *RAD27* at the lowest vs highest levels, with the highest rates produced by the highest expression levels. Stochastic variations in Rad27 or Rad52 expression seem to be mainly responsible for variation in HR rate, but other sources of gene expression heterogeneity probably amplify this phenomenon at the whole-population scale. However, we exclude that DNA damage heterogeneity is responsible for it because cells sorted at equal level of a DNA damage response protein (Bmh1) also harboured Rad27-dependent heterogeneity in HR rate. Moreover, viability is not more decreased by the phleomycin treatment in Rad27-high cells compared to the Rad27-low cells, suggesting that there is no more DNA damage that could explain higher expression in these cells. Finally, it is very unlikely that higher levels of Rad52 or Rad27 are in this state because of more underlying DNA damage that induces expression of HR genes rather than because of stochastic expression fluctuations. The contribution of cell cycle stage seems also weak because strong variations in HR rate between subpopulations are not correlated to strong changes in cell cycle stage, even if other experiments could confirm this point, for instance by blocking cells either in G1 or in G2, sorting them according to the expression level and measuring induced HR.

The correlation was unexpected concerning *RAD27* because rad27Δ mutants showed increased HR in various studies (Johnson et al., 1995;Sommers et al., 1995). In fact, HR was found to be essential in rad27Δ mutants (Symington, 1998). However it was observed that overexpression of Rad27 makes yeast cells sensitive to hydroxyurea (HU), methyl methanesulfonate (MMS) and bleomycine (Duffy et al., 2016; Becker et al., 2018). Additionally, the study by Duffy *et al* shows that the number of Rad52 spots increase when rad27 is overexpressed. The study by Becker *et al* shows that Rad27 overexpression impedes replication fork progression and leads to an accumulation of cells in mid-S phase. Therefore it could be proposed that a high Rad27 level could generates DNA nicks or DSB that would induce an increase in HR frequency. Moreover previous results on chicken cells already suggested that Rad27 could facilitate HR by removing divergent sequences at DNA break ends (Kikuchi et al., 2005) making coherent the relationship we observed, even if it has also been shown as playing a role in limiting HR between short sequences in yeast (Negritto et al., 2001). Finally, it is worth noting that we tested phenotypic effects of gene expression noise providing limited quantitative variations from cell-to-cell unlike deletion experiments. Our results on Rad27 provide such example of molecular effects of weakly imbalanced protein levels that are the opposite of those resulting from simple deletion.

Apart from simple deletion (Yuen et al., 2007), expression variations of numerous genes are known to affect genome stability (Stirling et al., 2011; Ang et al., 2016; Duffy et al., 2016). As expected these genes are mainly involved in DNA damage response (e.g. DNA repair and recombination) and chromosome maintenance. In yeast, large scale screening revealed that many genes impact genome stability either when deleted (Yuen et al., 2007) or when differentially expressed (Stirling et al., 2011; Zhu et al., 2015; Duffy et al., 2016). Genetic events analyzed in these studies range from loss of a full mini-chromosome that measure chromosome instability (Yuen et al., 2007; Stirling et al., 2011; Zhu et al., 2015;Duffy et al., 2016) to loss of an endogenous locus (the mating type locus *MAT* on chromosome III for instance) that detect more limited genetic modifications (Yuen et al., 2007; Stirling et al., 2011; Duffy et al., 2016). These different types of measurements explain why genes impacting HR activity as *RAD52* and *RAD27* are not detected in the former case (Zhu et al., 2015), and observed in the latter (Yuen et al., 2007).

Two limitations can be highlighted about these works. First, genetic events resulting from multiple possible molecular mechanisms are detected, rendering impossible the quantitative analysis of a specific pathway in terms of event frequency. Loss of *URA3* inserted among the *FLO1* tandem repeats specifically detects limited deletions occurring between dispersed repeated DNA through SSA (Verstrepen et al., 2005), thus allowing this quantitative measurement of a specific pathway activity. Second, as mentioned in a recent study (Keren et al., 2016), these genome-wide libraries of knock-outs, reduction-of-function and overexpression delineate the effects of extreme expression levels that are typically far from wild-type expression: they do not reveal the dependence of phenotype on expression variations that occur in the vicinity of wild-type level. The authors of this study explored the relationship between gene expression and phenotype along a large expression spectrum with small increments to provide more information on the sensitivity of cellular properties to the expression levels. Unfortunately no gene involved in DNA repair or recombination was part of the study. A former study in *E. coli* modulated the expression of the mismatch repair protein MutL at multiple different cellular levels and revealed that the frequency of deletion-generating recombination is inversely related to the amount of MutL while mismatch repair activity is insensitive to fluctuations in MutL (Elez et al., 2007). Nevertheless in all cases phenotypic measurements were performed on whole populations harboring various mean expression levels, even if they were only slightly different. The present study takes a further step by allowing testing the degree of heterogeneity in genome stability in the range of “natural” or “physiological” stochastic variations of genes involved in DNA replication, repair and recombination.

HR can produce gene copy number variations (CNV) if the distance between the repeated sequences is relatively short (Hastings et al., 2009). Indeed, the fact that resection reaches both repeats so that the break is repaired by SSA is less likely when the distance separating the repeats increases (Hastings et al., 2009). More generally, SSA is responsible for repeat-mediated rearrangements (Bhargava et al., 2016) and HR globally contains the intrinsic capacity to modify genetic material through gene conversion and crossing over (Guirouilh-Barbat et al., 2014). Thus it was highly conceivable that noise in the expression of genes affecting HR activity produces variable capacity to evolve (evolvability) (Capp, 2010), as recently suggested for mutagenesis in *E. coli* (Uphoff et al., 2016).

Interestingly from an evolutionary viewpoint, we observed that HR rate scales non-linearly with Rad27 levels. If the relationship was linear, the total amount of HR would depend only on the averaged Rad27 expression. On the contrary this non-linearity implies that mean doubling Rad27 levels do not lead to a doubling of HR rate. Total amount of HR cannot be explained solely by the population- or time-averaged Rad27 expression and slight modifications of the Rad27 mean expression level in the population could generate high variation in the total amount of HR and allow its rapid tuning without the need of strong expression variations or mutant alleles. Moreover, modifying Rad27 expression noise, while keeping the average expression level the same, would have an effect on the total amount of HR. Such modifications of noise levels have be considered as another way to modify HR rate at the whole-population level apart from modifications of mean levels. This also suggests that noise levels in the expression of genes affecting genome stability could be under positive or negative selection. This direct influence of gene expression noise on the rate of appearance of genetic variations has to be considered in addition to, and independently of, recent observations showing that evolvability is dependent on the level of noise in the expression of genes affecting resistance in selective environments because it shapes mutational effects (Bodi et al., 2017).

Finally the human *RAD27* homolog *FEN1* (Singh et al., 2008) and *RAD52* (Lieberman et al., 2016), as well as many other genes involved in DNA replication, repair and recombination (Lahtz and Pfeifer, 2011; Chae et al., 2016), can be over- or under-expressed in human cancers thus producing genetic instability (Stratton et al., 2009). One can suggest that these expression variations are selected for along with the beneficial genetic alterations they have produced, the initial source of variations being gene expression noise (Capp, 2010). Moreover noise could be globally increased in cancer cells, with consequences on genome instability (Capp, 2005; 2010; 2017). Given the diverse influences of gene expression noise on genotype variations that this work and other recent works (Bodi et al., 2017) revealed, the idea to control the level of expression noise among cancer cells might allow limiting evolvability, and escape from therapy (Capp, 2012; Brock et al., 2015). The same idea could be applied to microbial populations in the aim to stabilize production phenotypes for instance by avoiding the appearance of extreme subpopulations with high genome instability that would more probably lose interesting production features. Finally, this interplay between the genetic, epigenetic, and gene expression variabilities is a highly exciting field of investigation, and could help elucidating the degree to which noise levels are indeed under selection and the environmental conditions favoring such selection (Keren et al., 2016), especially when affecting genome stability. In conclusion, the present study revealed that gene expression variability can produce heterogeneous evolvability through homologous recombination from cell-to-cell, with probable consequences for instance in terms of stress response in microbial populations or evolution of cancel cell populations in oncogenesis and therapeutic response.

## Acknowledgements

This work was in part supported by the Agence Nationale de la Recherche (grant number ANR-12-JSV6-0006 to JPC). We are grateful to Delphine Lestrade and Julien Cescut from the Toulouse White Biotechnology consortium for flow cytometry facilities, to Adilia Dagkesamanskaia for her helpful contribution and to Sébastien Déjean for his advices on statistical analysis. We also thank Kevin J Verstrepen for providing the KV133 strain containing the recombination substrate. Finally we are thankful to Fayza Daboussi, Yvan Canitrot and Ivan Matic for critical reading of the manuscript.

## Author Contributions Statement

J.L., J.M.F and J.P.C. conceived and designed the experiments. J.L. performed experiments. J.L. and J.P.C. wrote the manuscript.

## References

Alexander, H.K., Mayer, S.I., and Bonhoeffer, S. (2017). Population Heterogeneity in Mutation Rate Increases the Frequency of Higher-Order Mutants and Reduces Long-Term Mutational Load. Mol Biol Evol 34, 419–436.

Alvaro, D., Lisby, M., and Rothstein, R. (2007). Genome-wide analysis of Rad52 foci reveals diverse mechanisms impacting recombination. PLoS Genet 3, e228.

Ang, J.S., Duffy, S., Segovia, R., Stirling, P.C., and Hieter, P. (2016). Dosage Mutator Genes in Saccharomyces cerevisiae: A Novel Mutator Mode-of-action of the Mph1 DNA Helicase. Genetics.

Balakrishnan, L., and Bambara, R.A. (2013). Flap endonuclease 1. Annu Rev Biochem 82, 119–138.

Bar-Even, A., Paulsson, J., Maheshri, N., Carmi, M., O’shea, E., Pilpel, Y., and Barkai, N. (2006). Noise in protein expression scales with natural protein abundance. Nat Genet 38, 636–643.

Becker, J.R., Gallo, D., Leung, W., Croissant, T., Thu, Y.M., Nguyen, H.D., Starr, T.K., Brown, G.W., and Bielinsky, A.K. (2018). Flap endonuclease overexpression drives genome instability and DNA damage hypersensitivity in a PCNA-dependent manner. Nucleic Acids Res 46, 5634–5650.

Bhargava, R., Onyango, D.O., and Stark, J.M. (2016). Regulation of Single-Strand Annealing and its Role in Genome Maintenance. Trends Genet 32, 566–575.

Blake, W.J., Balazsi, G., Kohanski, M.A., Isaacs, F.J., Murphy, K.F., Kuang, Y., Cantor, C.R., Walt, D.R., and Collins, J.J. (2006). Phenotypic consequences of promoter-mediated transcriptional noise. Mol Cell 24, 853–865.

Blake, W.J., M, K.A., Cantor, C.R., and Collins, J.J. (2003). Noise in eukaryotic gene expression. Nature 422, 633–637.

Bodi, Z., Farkas, Z., Nevozhay, D., Kalapis, D., Lazar, V., Csorgo, B., Nyerges, A., Szamecz, B., Fekete, G., Papp, B., Araujo, H., Oliveira, J.L., Moura, G., Santos, M.a.S., Szekely, T., Jr., Balazsi, G., and Pal, C. (2017). Phenotypic heterogeneity promotes adaptive evolution. PLoS Biol 15, e2000644.

Brock, A., Krause, S., and Ingber, D.E. (2015). Control of cancer formation by intrinsic genetic noise and microenvironmental cues. Nat Rev Cancer 15, 499–509.

Capp, J.P. (2005). Stochastic gene expression, disruption of tissue averaging effects and cancer as a disease of development. Bioessays 27, 1277–1285.

Capp, J.P. (2010). Noise-driven heterogeneity in the rate of genetic-variant generation as a basis for evolvability. Genetics 185, 395–404.

Capp, J.P. (2012). Stochastic gene expression stabilization as a new therapeutic strategy for cancer. Bioessays 34, 170–173.

Capp, J.P. (2017). Tissue disruption increases stochastic gene expression thus producing tumors: Cancer initiation without driver mutation. Int J Cancer 140, 2408–2413.

Chae, Y.K., Anker, J.F., Carneiro, B.A., Chandra, S., Kaplan, J., Kalyan, A., Santa-Maria, C.A., Platanias, L.C., and Giles, F.J. (2016). Genomic landscape of DNA repair genes in cancer. Oncotarget 7, 23312–23321.

Debrauwere, H., Loeillet, S., Lin, W., Lopes, J., and Nicolas, A. (2001). Links between replication and recombination in Saccharomyces cerevisiae: a hypersensitive requirement for homologous recombination in the absence of Rad27 activity. Proc Natl Acad Sci U S A 98, 8263–8269.

Dornfeld, K.J., and Livingston, D.M. (1992). Plasmid recombination in a rad52 mutant of Saccharomyces cerevisiae. Genetics 131, 261–276.

Duffy, S., Fam, H.K., Wang, Y.K., Styles, E.B., Kim, J.H., Ang, J.S., Singh, T., Larionov, V., Shah, S.P., Andrews, B., Boerkoel, C.F., and Hieter, P. (2016). Overexpression screens identify conserved dosage chromosome instability genes in yeast and human cancer. Proc Natl Acad Sci U S A 113, 9967–9976.

Dumont, B.L., Broman, K.W., and Payseur, B.A. (2009). Variation in genomic recombination rates among heterogeneous stock mice. Genetics 182, 1345–1349.

Elez, M., Radman, M., and Matic, I. (2007). The frequency and structure of recombinant products is determined by the cellular level of MutL. Proc Natl Acad Sci U S A 104, 8935–8940.

Engels, K., Giannattasio, M., Muzi-Falconi, M., Lopes, M., and Ferrari, S. (2011). 14-3-3 Proteins regulate exonuclease 1-dependent processing of stalled replication forks. PLoS Genet 7, e1001367.

Fraser, D., and Kaern, M. (2009). A chance at survival: gene expression noise and phenotypic diversification strategies. Mol Microbiol 71, 1333–1340.

Guirouilh-Barbat, J., Lambert, S., Bertrand, P., and Lopez, B.S. (2014). Is homologous recombination really an error-free process? Front Genet 5, 175.

Hastings, P.J., Lupski, J.R., Rosenberg, S.M., and Ira, G. (2009). Mechanisms of change in gene copy number. Nat Rev Genet 10, 551–564.

Hou, Y., Fan, W., Yan, L., Li, R., Lian, Y., Huang, J., Li, J., Xu, L., Tang, F., Xie, X.S., and Qiao, J. (2013). Genome analyses of single human oocytes. Cell 155, 1492–1506.

Johnson, R.E., Kovvali, G.K., Prakash, L., and Prakash, S. (1995). Requirement of the yeast RTH1 5’ to 3’ exonuclease for the stability of simple repetitive DNA. Science 269, 238–240.

Kauppi, L., Jeffreys, A.J., and Keeney, S. (2004). Where the crossovers are: recombination distributions in mammals. Nat Rev Genet 5, 413–424.

Keren, L., Hausser, J., Lotan-Pompan, M., Vainberg Slutskin, I., Alisar, H., Kaminski, S., Weinberger, A., Alon, U., Milo, R., and Segal, E. (2016). Massively Parallel Interrogation of the Effects of Gene Expression Levels on Fitness. Cell 166, 1282– 1294 e1218.

Kikuchi, K., Taniguchi, Y., Hatanaka, A., Sonoda, E., Hochegger, H., Adachi, N., Matsuzaki, Y., Koyama, H., Van Gent, D.C., Jasin, M., and Takeda, S. (2005). Fen-1 facilitates homologous recombination by removing divergent sequences at DNA break ends. Mol Cell Biol 25, 6948–6955.

Kim, D.R., Gidvani, R.D., Ingalls, B.P., Duncker, B.P., and Mcconkey, B.J. (2011). Differential chromatin proteomics of the MMS-induced DNA damage response in yeast. Proteome Sci 9, 62.

Kokoska, R.J., Stefanovic, L., Tran, H.T., Resnick, M.A., Gordenin, D.A., and Petes, T.D. (1998). Destabilization of yeast micro-and minisatellite DNA sequences by mutations affecting a nuclease involved in Okazaki fragment processing (rad27) and DNA polymerase delta (pol3-t). Mol Cell Biol 18, 2779–2788.

Kumar, R. (2017). An account of fungal 14-3-3 proteins. Eur J Cell Biol 96, 206–217.

Lahtz, C., and Pfeifer, G.P. (2011). Epigenetic changes of DNA repair genes in cancer. J Mol Cell Biol 3, 51–58.

Lieberman, R., Xiong, D., James, M., Han, Y., Amos, C.I., Wang, L., and You, M. (2016). Functional characterization of RAD52 as a lung cancer susceptibility gene in the 12p13.33 locus. Mol Carcinog 55, 953–963.

Liu, J., Francois, J.M., and Capp, J.P. (2016). Use of noise in gene expression as an experimental parameter to test phenotypic effects. Yeast 33, 209–216.

Liu, J., Martin-Yken, H., Bigey, F., Dequin, S., Francois, J.M., and Capp, J.P. (2015). Natural yeast promoter variants reveal epistasis in the generation of transcriptional-mediated noise and its potential benefit in stressful conditions. Genome Biol Evol 7, 969–984.

Lu, S., Zong, C., Fan, W., Yang, M., Li, J., Chapman, A.R., Zhu, P., Hu, X., Xu, L., Yan, L., Bai, F., Qiao, J., Tang, F., Li, R., and Xie, X.S. (2012). Probing meiotic recombination and aneuploidy of single sperm cells by whole-genome sequencing. Science 338, 1627–1630.

Moore, C.W. (1988). Internucleosomal cleavage and chromosomal degradation by bleomycin and phleomycin in yeast. Cancer Res 48, 6837–6843.

Negritto, M.C., Qiu, J., Ratay, D.O., Shen, B., and Bailis, A.M. (2001). Novel function of Rad27 (FEN-1) in restricting short-sequence recombination. Mol Cell Biol 21, 2349–2358.

New, J.H., Sugiyama, T., Zaitseva, E., and Kowalczykowski, S.C. (1998). Rad52 protein stimulates DNA strand exchange by Rad51 and replication protein A. Nature 391, 407–410.

Newman, J.R., Ghaemmaghami, S., Ihmels, J., Breslow, D.K., Noble, M., Derisi, J.L., and Weissman, J.S. (2006). Single-cell proteomic analysis of S. cerevisiae reveals the architecture of biological noise. Nature 441, 840–846.

Paques, F., and Haber, J.E. (1999). Multiple pathways of recombination induced by double-strand breaks in Saccharomyces cerevisiae. Microbiol Mol Biol Rev 63, 349–404.

Raser, J.M., and O’shea, E.K. (2005). Noise in gene expression: origins, consequences, and control. Science 309, 2010–2013.

Rattray, A.J., and Symington, L.S. (1994). Use of a chromosomal inverted repeat to demonstrate that the RAD51 and RAD52 genes of Saccharomyces cerevisiae have different roles in mitotic recombination. Genetics 138, 587–595.

Silander, O.K., Nikolic, N., Zaslaver, A., Bren, A., Kikoin, I., Alon, U., and Ackermann, M. (2012). A genome-wide analysis of promoter-mediated phenotypic noise in Escherichia coli. PLoS Genet 8, e1002443.

Singh, P., Yang, M., Dai, H., Yu, D., Huang, Q., Tan, W., Kernstine, K.H., Lin, D., and Shen, B. (2008). Overexpression and hypomethylation of flap endonuclease 1 gene in breast and other cancers. Mol Cancer Res 6, 1710–1717.

Skotheim, J.M., Di Talia, S., Siggia, E.D., and Cross, F.R. (2008). Positive feedback of G1 cyclins ensures coherent cell cycle entry. Nature 454, 291–296.

Smith, M.C., Sumner, E.R., and Avery, S.V. (2007). Glutathione and Gts1p drive beneficial variability in the cadmium resistances of individual yeast cells. Mol Microbiol 66, 699–712.

Sommers, C.H., Miller, E.J., Dujon, B., Prakash, S., and Prakash, L. (1995). Conditional lethality of null mutations in RTH1 that encodes the yeast counterpart of a mammalian 5’-to 3’-exonuclease required for lagging strand DNA synthesis in reconstituted systems. J Biol Chem 270, 4193–4196.

Song, B., and Sung, P. (2000). Functional interactions among yeast Rad51 recombinase, Rad52 mediator, and replication protein A in DNA strand exchange. J Biol Chem 275, 15895–15904.

Stirling, P.C., Bloom, M.S., Solanki-Patil, T., Smith, S., Sipahimalani, P., Li, Z., Kofoed, M., Ben-Aroya, S., Myung, K., and Hieter, P. (2011). The complete spectrum of yeast chromosome instability genes identifies candidate CIN cancer genes and functional roles for ASTRA complex components. PLoS Genet 7, e1002057.

Stratton, M.R., Campbell, P.J., and Futreal, P.A. (2009). The cancer genome. Nature 458, 719–724.

Symington, L.S. (1998). Homologous recombination is required for the viability of rad27 mutants. Nucleic Acids Res 26, 5589–5595.

Symington, L.S. (2002). Role of RAD52 epistasis group genes in homologous recombination and double-strand break repair. Microbiol Mol Biol Rev 66, 630-670, table of contents.

Symington, L.S., Rothstein, R., and Lisby, M. (2014). Mechanisms and regulation of mitotic recombination in Saccharomyces cerevisiae. Genetics 198, 795–835.

Thomas, B.J., and Rothstein, R. (1989). Elevated recombination rates in transcriptionally active DNA. Cell 56, 619–630.

Tishkoff, D.X., Filosi, N., Gaida, G.M., and Kolodner, R.D. (1997). A novel mutation avoidance mechanism dependent on S. cerevisiae RAD27 is distinct from DNA mismatch repair. Cell 88, 253–263.

Uphoff, S., Lord, N.D., Okumus, B., Potvin-Trottier, L., Sherratt, D.J., and Paulsson, J. (2016). Stochastic activation of a DNA damage response causes cell-to-cell mutation rate variation. Science 351, 1094–1097.

Usui, T., and Petrini, J.H. (2007). The Saccharomyces cerevisiae 14-3-3 proteins Bmh1 and Bmh2 directly influence the DNA damage-dependent functions of Rad53. Proc Natl Acad Sci U S A 104, 2797–2802.

Verstrepen, K.J., Jansen, A., Lewitter, F., and Fink, G.R. (2005). Intragenic tandem repeats generate functional variability. Nat Genet 37, 986–990.

Viney, M., and Reece, S.E. (2013). Adaptive noise. Proc Biol Sci 280, 20131104.

Wang, J., Fan, H.C., Behr, B., and Quake, S.R. (2012). Genome-wide single-cell analysis of recombination activity and de novo mutation rates in human sperm. Cell 150, 402–412.

Yuen, K.W., Warren, C.D., Chen, O., Kwok, T., Hieter, P., and Spencer, F.A. (2007). Systematic genome instability screens in yeast and their potential relevance to cancer. Proc Natl Acad Sci U S A 104, 3925–3930.

Zheng, L., and Shen, B. (2011). Okazaki fragment maturation: nucleases take centre stage. J Mol Cell Biol 3, 23–30.

Zhu, J., Heinecke, D., Mulla, W.A., Bradford, W.D., Rubinstein, B., Box, A., Haug, J.S., and Li, R. (2015). Single-Cell Based Quantitative Assay of Chromosome Transmission Fidelity. G3 (Bethesda) 5, 1043–1056.

Zopf, C.J., Quinn, K., Zeidman, J., and Maheshri, N. (2013). Cell-cycle dependence of transcription dominates noise in gene expression. PLoS Comput Biol 9, e1003161.

