## Supplementary Material for "Gene expression noise produces cell-to-cell heterogeneity in eukaryotic homologous recombination rate"

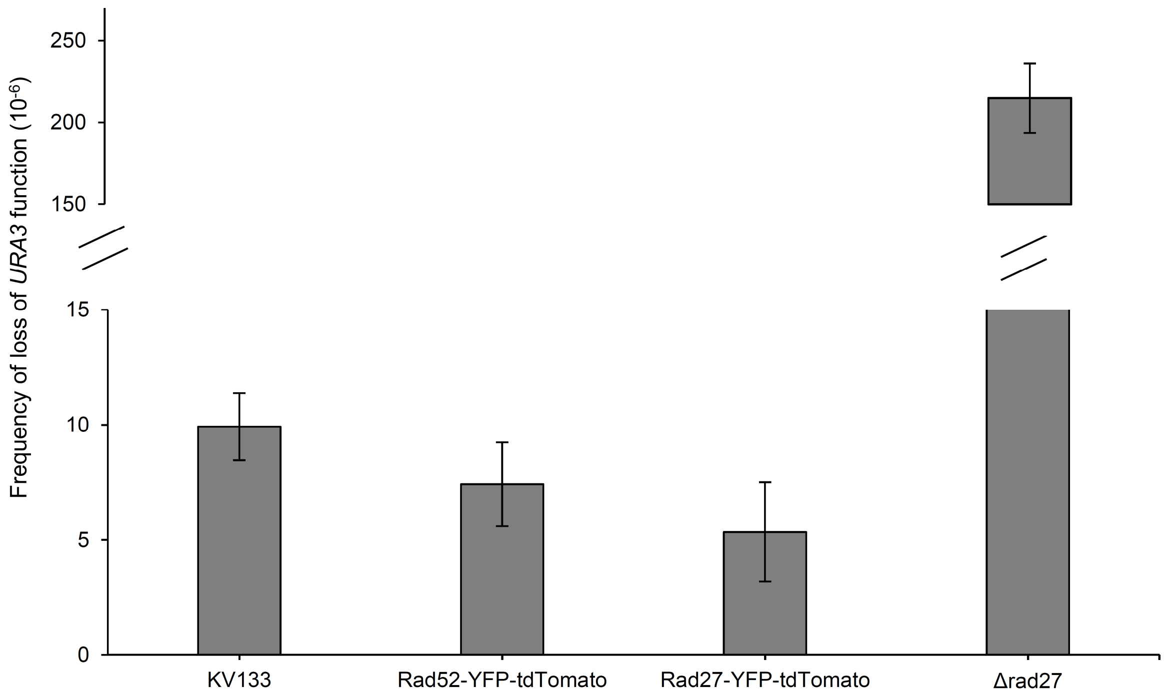


**Supplementary Figure 1.** Average spontaneous frequency of loss of *URA3* function in whole populations. Results are the mean of 3 independent experiments with standard deviation.

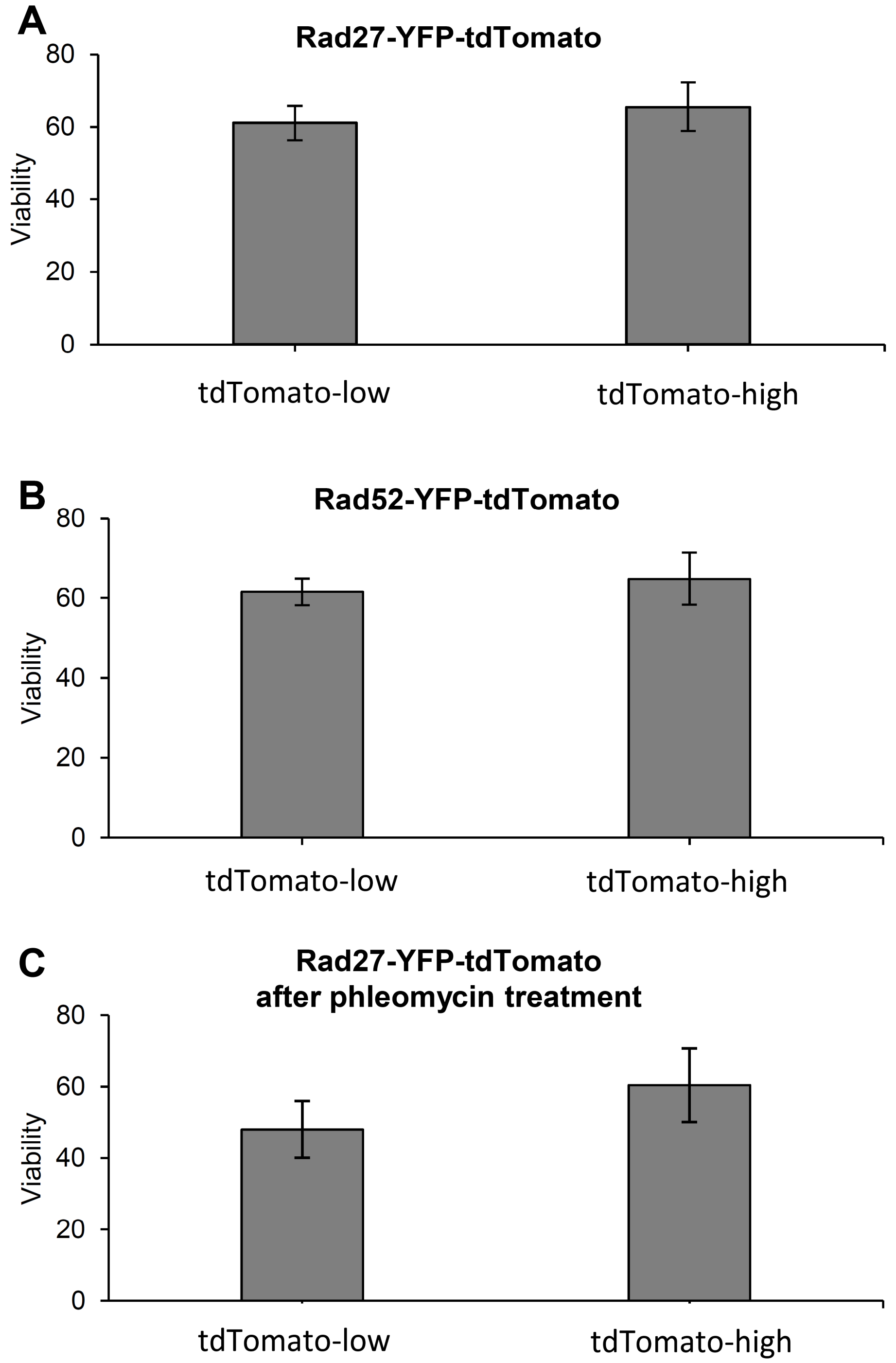


**Supplementary Figure 2.** Viability in the different subpopulations used to measure frequencies of loss of *URA3* function in Figure 2. Results are the mean of 3 independent experiments with standard deviation.

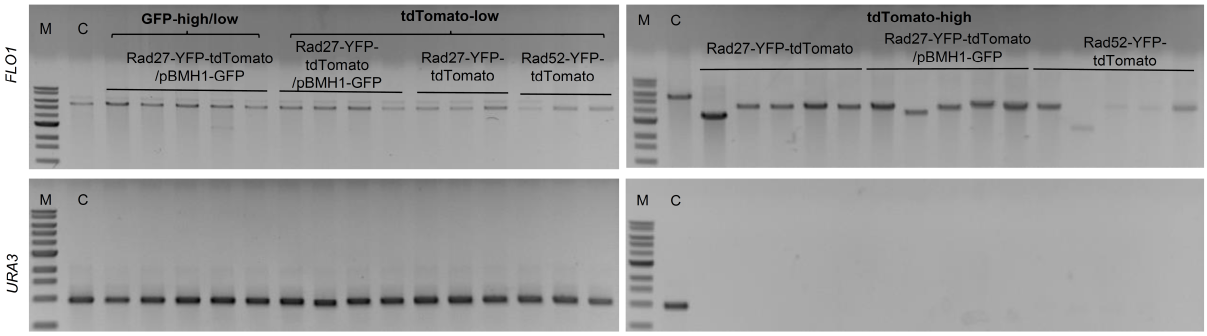


**Supplementary Figure 3**. PCR amplification of the *FLO1* and *URA3* alleles in different clones obtained on 5-FOA plates. Examples of PCR amplification of the new *FLO1* alleles in 5-FOA resistant clones showing that their length is modified in the high-expressing subpopulations, and not in the low-expressing subpopulations compared to the control strain (C). PCR amplification of the *URA3* gene in the same clones showed that it is lost by HR in the high-expressing subpopulations and still present in the low-expressing subpopulations.

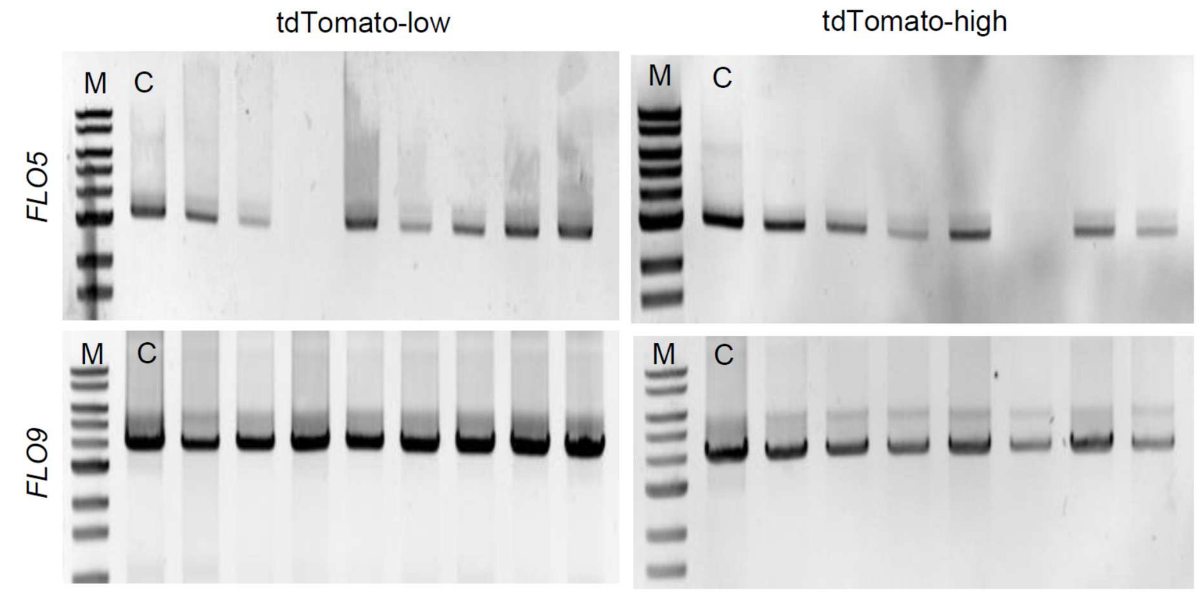


**Supplementary Figure 4**. PCR amplification of the *FLO5* and *FLO9* alleles in different clones obtained on 5-FOA plates. Examples of PCR amplification of other loci containing tandem repeats (*FLO5* and *FLO9*) in the *FLO1* recombinant clones compared to the control strain (C).

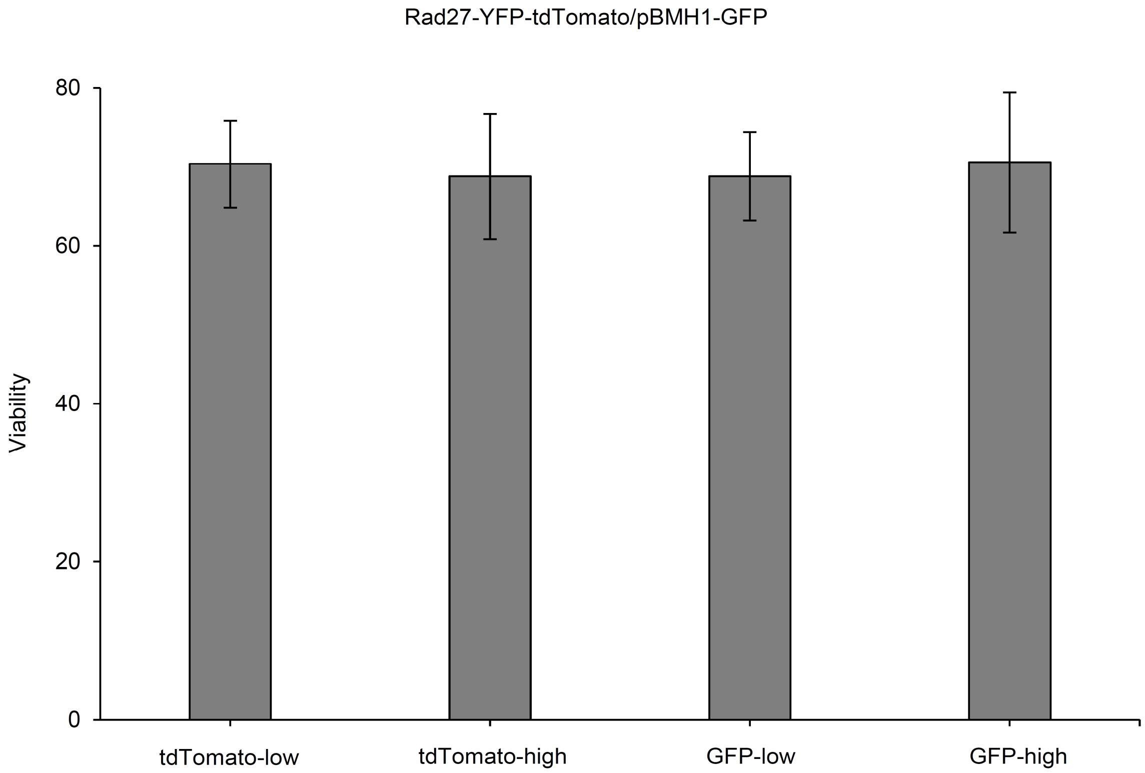


**Supplementary Figure 5**. Viability in the different subpopulations used to measure frequencies of loss of *URA3* function in Figure 4. Results are the mean of 3 independent experiments with standard deviation.

**Supplementary Table 1**. Raw data of the recombination rate analyses.

See enclosed .xlsx file

| Name | Gene type | Reference/ Source |
|---|---|---|
| KV133 | BY4742 (MATα; his3Δ1; leu2Δ0; lys2Δ0; ura3Δ0) FLO1::URA3 (URA3 inserted in the middle of the tandem repeats) | Verstrepen KJ *et al.* , Intragenic tandem repeats generate functional variability. Nat Genet 37, 986-990 (2005) |
| JA0200 | KV133 *LEU2* | This study |
| JA0217 | JA0200 *Δrad27::LYS2* | This study |
| JA0219 | JA0200 *RAD27 -YFP -kanR* | This study |
| JA0220 | JA0200 *RAD52-YFP-kanR* | This study |
| JA0240 | JA0200 *RAD27-YFP-tdTomato-SpHis5* | This study |
| JA0241 | JA0200 *RAD52-YFP-tdTomato-SpHis5* | This study |
| JA0242 | JA0240 Δ*leu2::pBMH1-yEGFP* | This study |
| JA0243 | JA0200 Δ*leu2::pBMH1-yEGFP* | This study |

**Supplementary Table 2**. List of the strains used in this study

| Name in the article | Name in the stock | Sequence |
|---|---|---|
| F1 | LEU2-for | ATGACAAAACCTCTTCCGAT |
| R1 | LEU2-rev | CCCTCCTCCTTGTCAATATT |
| C1 | CHECK-4gene-for | CTCAACATAACGAGAACACACA |
| C2 | CHECK-4gene-plus-rev | AATGGTCAGGTCATTGAGTG |
| F2 | Rad27-add-YFP-kan-for | AAAATAAAAAATTGAACAAAAATAAGAATAAAGTCACAAAGGGAAGAAGACGGATCCCCGGGTTAATTAA |
| R2 | Rad27-YFP-Add-kan-rev | AGGTAAGAATGAAAAATTCCACGTTCAAGTTCCCAGAAAAACTGGCAAAATACTTTCTGCGCACTTAACT |
| C3 | Check-Rad27-YFP-for | GCCACCAAAGGAGAAGGAACTT |
| C4 | Check-Rad-YFP-rev | TTGGGATCTTTCGAAAGGGC |
| F3 | Rad52- add-YFP-kan-for | GAGAAGTTGGAAGACCAAAGATCAATCCCCTGCATGCACGCAAGCCTACTCGGATCCCCGGGTTAATTAA |
| R3 | Rad52-YFP-Add-kan-rev | CTTGTAAAATAATAAGAATTTTTTATTCGATTTAAAGTAAATATTAATACTACTTTCTGCGCACTTAACT |
| C5 | Check-Rad52-YFP –for | CGCGAGGGATTCTGTCTATGAA |
| F4 | AdtdToma-YFP-for | TGTTACTGCTGCTGGTATTACCCATGGTATGGATGAATTGTACAAAATTAACATGGTGAGCAAGGG |
| R4 | AdtdToma-rev | AAATGACAAGTTCTTGAAAACAAGAATCTTTTTATTGTCAGTACTTTACTTGTACAGCTCGTCCATG |
| C6 | che-tdTomo-rev | CCCTCGATCTCGAACTCGTG |
| F5 | FLO1-F3 | ATCGCTATATGTTTTTGGCAGTCTTTA |
| R5 | FLO1-5-9-R4 | TTAAATAATTGCCAGCAATAAGGACG |
| F6 | Lys2-Rad27-For | CGTTGACAGCATACATTGGAAAGAAATAGGAAACGGACACCGGAAG CTTCTGCTGCGTATTATTCTGC |
| R6 | Lys2-Rad27-Rev | TGCCAAGGTGAAGGACCAAAAGAAGAAAGTGGAAAAAGAACCCCCTTAAGCTGCTGCGGAGCTT |
| C7 | CHECK-Lys2-For-rad27 | CCGGCTGGTAAGTTATGATAGA |
| C8 | CHECK-Lys2-Rev | GCATTGTCCTGGAAAATGTC |
| C9 | CHECK-PJL2-for | ACATACATAAACATACGCGC |
| C10 | CHECK-URA-rev | TTAGTTTTGCTGGCCGCATC |
| C11 | URA3-F1 | GATTCGGTAATCTCCGAGCAGA |
| C12 | FLO9-F2 | TTATTGTTTACTACTAGCCATCGTCACA |
| C13 | FLO5-F3 | GCACACCACTGCATATTTTTGGTAA |

**Supplementary Table 3**. List of the primers used in this study
